# The McdAB system positions α-carboxysomes in proteobacteria

**DOI:** 10.1101/2020.08.11.246918

**Authors:** Joshua S. MacCready, Lisa Tran, Joseph L. Basalla, Pusparanee Hakim, Anthony G. Vecchiarelli

## Abstract

Carboxysomes are protein-based organelles essential for carbon fixation in cyanobacteria and proteobacteria. Previously, we showed that the cyanobacterial nucleoid is utilized as a surface for the equidistant-spacing of β-carboxysomes across cell lengths by a two-component system (McdAB) in the model cyanobacterium *Synechococcus elongatus* PCC 7942. More recently, we found that McdAB systems are widespread among β-cyanobacteria, which possess β-carboxysomes, but are absent in α-cyanobacteria, which possess structurally distinct α-carboxysomes. Since cyanobacterial α-carboxysomes are thought to have arisen in proteobacteria and were subsequently horizontally transferred into cyanobacteria, this raised the question whether α-carboxysome containing proteobacteria possess a McdAB system for positioning α-carboxysomes. Here, using the model chemoautotrophic proteobacterium *H. neapolitanus*, we show that a McdAB system distinct from that of β-cyanobacteria operates to position α-carboxysomes across cell lengths. We further show that this system is widespread among α-carboxysome containing proteobacteria and that cyanobacteria likely inherited an α-carboxysome operon from a proteobacterium lacking the *mcdAB* locus. These results demonstrate that McdAB is a cross-phylum two-component system necessary for positioning α- and β-carboxysomes. The findings have further implications for understanding the positioning of other bacterial protein-based organelles involved in diverse metabolic processes.

## Introduction

Bacterial Microcompartments (BMCs) are a family of large cytosolic protein-based organelles that encapsulate sensitive metabolic processes to provide microbes with a distinct environmental growth advantage **(Reviewed in: Kerfeld et al., 2018)**. Diverse in structure and function, BMCs have been identified across 23 bacterial phyla and ~20% of all sequenced bacterial genomes **(Axen et al., 2014)**, so their functions are of great ecological, evolutionary, biotechnological and medical interest. The model BMC is the carboxysome, which is named for its involvement in carbon-fixation. Carboxysomes are classified as either α or β depending on the type of Ribulose-1,5-bisphosphate carboxylase/oxygenase (RuBisCO) they encapsulate; α-carboxysomes contain RuBisCO form 1A, β-carboxysomes contain RuBisCO form 1B, and both carboxysome types possess a distinct set of protein components necessary for encapsulation **(Figure 1AB) (Reviewed in: Rae et al., 2013)**. Since RuBisCO can utilize either CO_2_ or O_2_ as a substrate, driving either the Calvin-Benson-Bassham cycle (CO_2_) or the wasteful process of photorespiration (O_2_), encapsulation of RuBiscO and carbonic anhydrase within a selectively permeable protein shell generates a high internal CO_2_ environment within carboxysomes that minimizes photorespiration **(Price and Badger, 1989; Lieman-Hurwitz et al., 1991; Badger et al., 1998; Tcherkez et al., 2006)**. Through this mechanism, carboxysomes contribute to greater than 35% of global carbon fixation through atmospheric CO2 assimilation **(Dworkin, 2006; Kerfeld and Melnicki, 2016)**.

**Figure 1:**
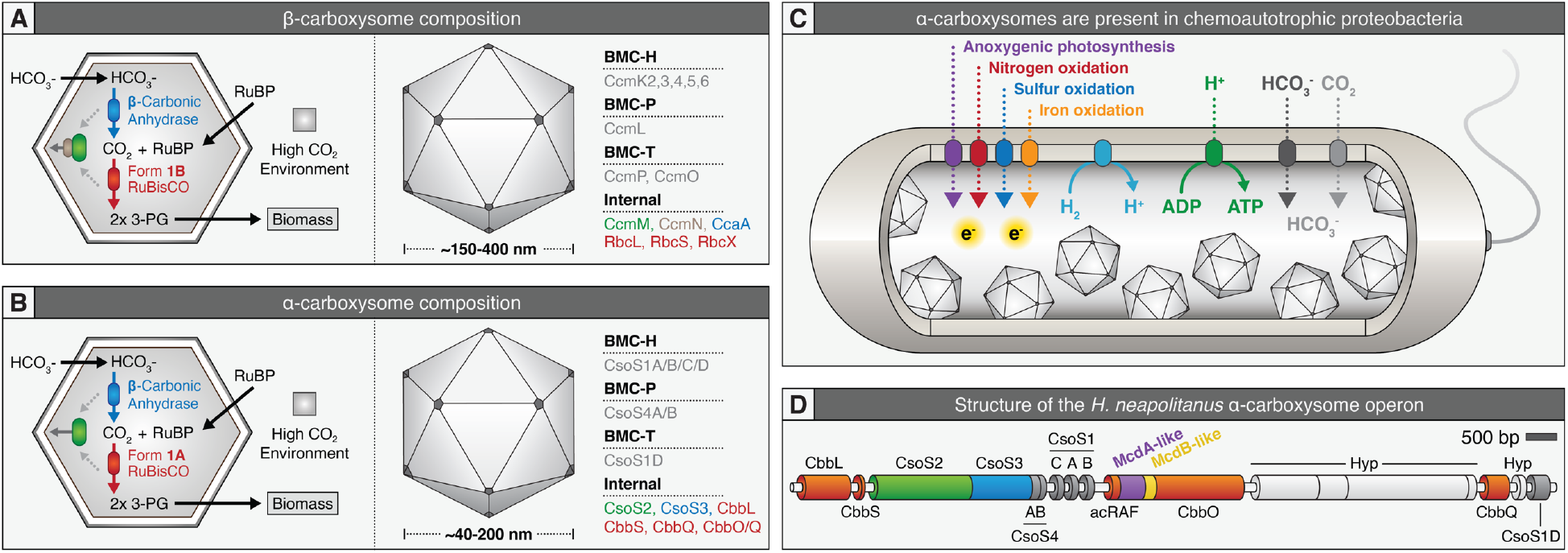
Proteobacterial *cso* operons contain McdAB-like sequences. **(A)** Cartoon illustration of internal reactions (left) and overall known components (right) of β-carboxysomes and **(B)** α-carboxysomes. **(C)** α-carboxysome containing proteobacteria display diverse metabolic capacities. (D) Genetic arrangement of the *H. neapolitanus* α-carboxysome operon. Red = RuBisCO or RuBisCO associated, green = CsoS2 which mediates RuBisCO/shell interaction, blue = carbonic anhydrase, purple = putative *mcdA* gene, yellow = putative *mcdB* gene, dark grey = shell component, light grey = hypothetical protein (Hyp).

In our previous study, we identified and characterized the two-component system McdAB (<u>M</u>aintenance of <u>C</u>arboxysome <u>D</u>istribution protein <u>A</u> and <u>B</u>) required for maintaining the equidistant positioning of multiple carboxysomes along the nucleoid in the model rod-shaped cyanobacterium *Synechococcus elongatus* PCC 7942 **(MacCready et al., 2018)**. We found that McdA, a ParAtype ATPase, non-specifically bound DNA in the presence of ATP. McdB, a novel small disordered protein, localized to carboxysomes and stimulated the ATPase activity of McdA, removing McdA from the nucleoid. Once removed, McdA then recycled its nucleotide and rebound the nucleoid at locations with the lowest concentration of McdB. The local removal of McdA created depletion zones in the vicinity of McdB-bound carboxysomes that subsequently evolved into emergent global oscillations of McdA. McdB-bound carboxysomes therefore generate and exploit dynamic McdA gradients to move in a directed and persistent manner towards increased concentrations of McdA on the nucleoid. Since we found that carboxysomes clustered in the absence of the McdAB system, we concluded that its Brownian-ratchet based distribution mechanism functioned primarily as an “anti-aggregation” system while simultaneously ensuring proper inheritance of carboxysomes.

More recently, we found that two distinct types of McdAB systems (Type 1 and Type 2) exist among β-cyanobacteria, which possess β-carboxysomes **(MacCready et al., 2019)**. Surprisingly however, we also found that α-cyanobacteria, which possess α-carboxysomes, completely lack the McdAB system. Despite this observation, we previously reported that many chemoautotrophic and anoxygenic photosynthetic proteobacteria, which derive electrons from molecular hydrogen or reduced nitrogen, sulfur, and metal compounds **(Figure 1C)**, also possess α-carboxysomes and potentially encoded for McdA and McdB within their carboxysome (*cso*) operon **(Figure 1D**) **(MacCready et al., 2018)**. Interestingly, α-carboxysomes are thought to have arisen in proteobacteria and were subsequently horizontally transferred into cyanobacteria, resulting in the distinct α-cyanobacterial lineage. Thus, it remains unclear whether proteobacterial α-carboxysomes are positioned by a McdAB system and whether this system evolved prior to or after horizontal transfer into cyanobacteria.

Here, we show that proteobacteria possess a novel McdAB system, which we term α-McdAB, to position α-carboxysomes. In the sulfur-oxidizing chemoautotrophic proteobacterium *Halothiobacillus neapolitanus* c2, we show that uniform distributions of α-carboxysomes are lost in the absence of either α-McdA or α-McdB. Similar to the β-McdAB system in *S. elongatus*, we found that *H. neapolitanus* α-McdA and α-McdB interact and that α-McdB co-localizes with α-carboxysomes *in vivo*. Furthermore, we found that the α-McdAB system is widespread among α-carboxysome containing proteobacteria, many possessing two distinct copies of α-McdB. Lastly, we determined the oligomeric state of McdB representatives from the three major known types and found that β-McdB Type 1 forms a hexamer, β-McdB Type 2 forms a dimer, and α-McdB remains a monomer; results which further our understanding of how different McdB proteins interface with their cognate McdA and carboxysome cargo. Collectively, these results reveal a widespread McdAB system required for positioning structurally distinct α-carboxysomes in proteobacteria, provide insights into the evolution of α-carboxysomes between cyanobacteria and proteobacteria, and have broader implications for understanding the subcellular organization of BMCs outside of carboxysomes.

## Results and Discussion

### McdA and McdB maintain carboxysome distributions in *H. neapolitanus*

We began this study by testing the hypothesis that the McdA-like coding sequence that we previously identified within proteobacterial *cso* operons functioned to maintain their α-carboxysome distributions **(See Figure 8C – MacCready et al., 2018)**. Likewise, while sharing no identifiable characteristics with known cyanobacterial McdB sequences, we also tested whether the small coding sequence following the McdA-like sequence was a functional homolog to cyanobacterial McdB. To explore this, we used the sulfur-oxidizing chemoautotroph *Halothiobacillus neapolitanus* c2 (hereafter *H. neapolitanus)*, which has been established as a model system for studying α-carboxysome biology in proteobacteria **(Shively et al., 1973; Cannon and Shively, 1983; Holthuijzen et al., 1987; Schmid et al., 2006; Dou et al., 2008; Rae et al., 2013; Sutter et al., 2015; Oltrogge et al., 2020)**. First, in *H. neapolitanus*, we fused the native gene that encodes for the small subunit of RuBisCO (CbbS) with the fluorescent protein mTurquoise2 to form CbbS-mTQ **(Goedhart et al., 2012)**. Here, our CbbS-mTQ signal appeared as several dynamic but overlapping foci **(Video 1)**, which from a single image, looks like a nearly homogeneous signal across the length of individual *H. neapolitanus* cells **(Figure 2A)**. In *S. elongatus*, fluorescent-labeled carboxysomes are sufficiently separated from one another to be resolved **(Savage et al., 2010; Cameron et al., 2013; MacCready et al., 2018)**. In general, *H. neapolitanus* carboxysomes are greater in number (4 to 18) and smaller (40 to 200 nm diameter) compared to the fewer (3 to 5) and larger (150 to 300 nm) β-carboxysomes of *S. elongatus* **(Cannon et al., 2001; Schmid et al., 2006; Savage et al., 2010; Rae et al., 2013; MacCready et al., 2018; Sun et al., 2019)**. Therefore, we interpret this data as α-carboxysomes being distributed across the cell length **(Figure 2A)**. But the smaller size of α-carboxysomes combined with their high-copy number precludes single carboxysome resolution using traditional fluorescence microscopy imaging of *H. neapolitanus* cells, which are also relatively small (0.5 x 1.5 μm). We therefore performed Transmission Electron Microscopy (TEM) to determine if this CbbS-mTQ signal in wild-type cells indeed represented assembled and distributed carboxysomes. Consistent with our interpretation of the fluorescent imaging, we frequently found that α-carboxysomes were distributed over the nucleoid region of the cell and down the medial axis **Figure 2B)**.

**Figure 2:**
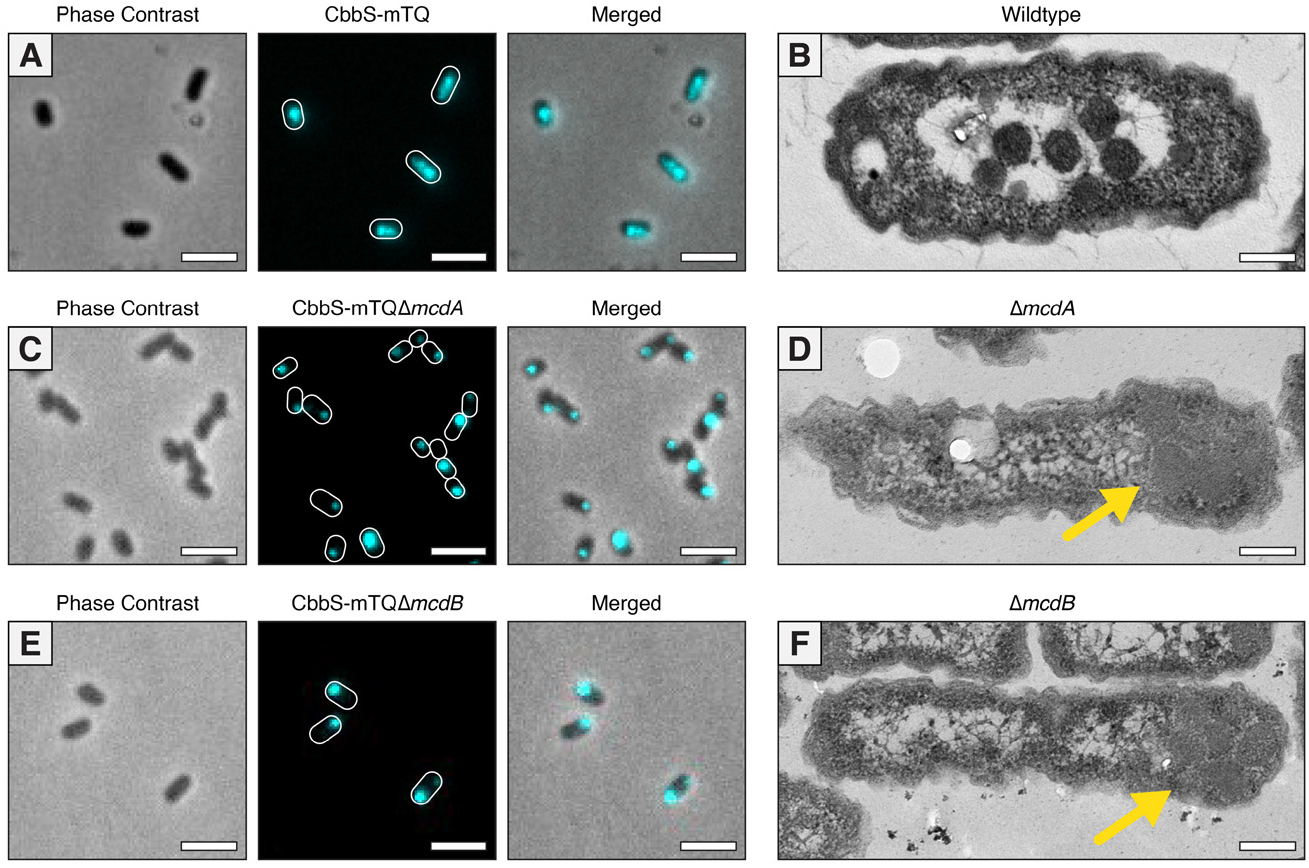
McdA and McdB position α-carboxysomes. **(A)** Native CbbS-mTQ signal is distributed across cell lengths. Scale bar = 2 μm. (B) α-carboxysomes are confined to the nucleoid in WT cells. Scale bar = 200 nm. (C) Homogenous distribution of α-carboxysomes is lost in the absence of McdA. ~76% of n=210 cells had a single polar focus and 7% displayed two foci at opposing poles. ~17% of cells lacked a focus. Scale bar = 2 μm. (D) Polar localization of assembled α-carboxysomes in the absence of McdA (yellow arrow). Scale bar = 200 nm. (E) Homogenous distribution of α-carboxysomes is lost in the absence of McdB. ~66% of n=220 cells had a single polar focus and 5% displayed two foci at opposing poles. ~29% of cells lacked a focus. Scale bar = 2 μm. (F) Polar localization of assembled α-carboxysomes in the absence of McdB (yellow arrow). Scale bar = 200 nm.

Next, using this CbbS-mTQ α-carboxysome reporter, we individually deleted our candidate *mcdA* or *mcdB* genes. Confirming our hypothesis, in the absence of either *mcdA* or *mcdB*, we found that the homogenous distribution of α-carboxysomes down the cell length was lost in either mutant strain **(Figure 2CE)**. Instead, the majority of cells had a single high-intensity CbbS-mTQ focus at one pole (~76% of n=210 cells for *ΔmcdA* and ~66% of n=220 cells for Δ*mcdB*). Additionally, 7% of *ΔmcdA* and 5% of *ΔmcdB* cells displayed two foci at opposing poles. The remaining cells (17% of *ΔmcdA* and 29% of Δ*mcdB*) lacked a CbbS-mTQ focus entirely, suggesting that these cells did not inherit α-carboxysomes after division and have yet to assemble them *de novo*. We performed TEM on Δ*mcdA* and Δ*mcdB* strains to determine if these massive foci of CbbS-mTQ represented assembled carboxysomes or amorphous aggregates. In the absence of either *mcdA* or *mcdB*, the vast majority of cells (~75% of n=214 cells) had a single cluster of multiple assembled carboxysomes **(Figure 1DF)**. Once again, the TEM confirmed our interpretation of the fluorescent imaging. Now that we have identified the McdAB systems of α and β carboxysomes, from this point forward, we distinguish the two systems according to the type of carboxysome being distributed: α-McdAB systems position proteobacterial α-carboxysomes and β-McdAB systems position cyanobacterial β-carboxysomes.

These findings are intriguing on multiple fronts. First, α-carboxysomes are typically much smaller and more numerous than β-carboxysomes in cells **(Cannon et al., 2001; Schmid et al., 2006; Rae et al., 2013; Sun et al., 2019)**. Therefore, given that *H. neapolitanus* cells are also smaller (0.5 x 1.5 microns) than *S. elongatus* cells (1.3 x 3 microns), it could have been reasoned that α-carboxysomes would rely on random diffusion for their inheritance and distribution in the cell; whereas the larger size and lower copy number of cyanobacterial β-carboxysomes necessitate an active positioning system. However, since we found that assembled α-carboxysomes aggregated towards the polar region of cells in the absence of α-McdA or α-McdB, reminiscent of β-carboxysomes in the absence of β-McdA or β-McdB **(MacCready et al., 2018)**, we conclude that all McdAB systems actively position carboxysomes to not only ensure proper inheritance following cell division, but to primarily function as an anti-aggregation system. Consistently, extreme α- and β-carboxysome aggregation has also been observed when heterologously expressed in plant chloroplasts in the absence of their cognate McdAB systems **(Lin et al., 2014; Long et al., 2018).**

In addition to α- and β-carboxysome size and quantity being significantly different between *S. elongatus* and *H. neapolitanus*, so is their nucleoid biology. While *H. neapolitanus* has one or two copies of its chromosome, *S. elongatus* is polypoid; harboring as many as 10 copies. Some β-cyanobacteria that we have shown to encode an McdAB system can have over 200 chromosome copies **(Griese et al., 2011; Watanabe, 2020)**. Since McdA uses the nucleoid as a matrix for positioning McdB-bound carboxysomes, how these significant differences in chromosome content influence McdA dynamics and carboxysome positioning is an outstanding question. It is possible that, like carboxysome and chromosome content **(Ohbayashi et al., 2019; Sun et al., 2019)**, McdAB protein levels may also be regulated by environmental growth conditions, such as nutrient availability (e.g., phosphate or nitrogen). Together, these findings are consistent with our previous characterization of the McdAB in positioning β-carboxysomes in the cyanobacterium *S. elongatus* and expand our understanding of α-carboxysome biology **(MacCready et al., 2018)**.

What is the physiological function of carboxysome positioning? While α- and β-carboxysome aggregation does not result in a high CO_2_-requiring phenotype **(MacCready et al., 2018; Desmarais et al., 2019)**, we recently found that β-carboxysome aggregation in *S. elongatus* does result in slower cell growth, cell elongation, and asymmetric cell division; presumably due to decreased carbon-fixation by RuBisCO in these aggregates **(Rillema et al., 2020)**. Moreover, it was recently found that degradation of inactive carboxysomes occurs at polar regions of a cyanobacterial cell **(Hill et al., 2020)**. How the polar localization of carboxysome aggregates, from a lack of a McdAB system, influences carboxysome function and turnover remains unclear. Therefore, the need to study α- and β-McdAB systems under varying biotic and abiotic conditions is warranted to fully understand the impact of carboxysome mispositioning and aggregation on cellular physiology.

### Without the McdAB system, clustered carboxysomes are nucleoid excluded to the cell poles

Macromolecular crowding of the cytoplasm in combination with the nucleoid acting as a formidable diffusion barrier have been shown to be responsible for the polar localization of several large intracellular bodies in bacteria **(Straight et al., 2007, Winkler et al., 2010)**, such as plasmids lacking an active positioning system **(Erdmann et al., 1999; Ringgaard et al., 2009)**. Without a ParA-system, plasmids that were once distributed equally along the nucleoid length, become nucleoid ‘excluded’ – the passive effect of the nucleoid as a diffusion barrier to mesoscale bodies in the cytoplasm **(Planchenault et al., 2020)**. Therefore, it seems that ParA-based positioning systems, such as McdAB, not only overcome the nucleoid as a diffusion barrier, but exploit it as a matrix for cargo movement and positioning.

*H. neapolitanus* is a halophile capable of growing under extreme hyperosmotic conditions where the cell membrane pushes against the compacted nucleoid, forming a barrier in the cell that can even restrict the diffusion of GFP **(Van Den Bogaart et al., 2007; Konopka et al., 2009)**. We therefore asked if the polar localization of carboxysome clusters in the Δ*mcdA* and Δ*mcdB* strains is the result of nucleoid exclusion. To explore this question, we used Sytox Orange to stain the nucleoids of WT, Δ*α-mcdA*, and Δ*α-mcdB H. neapolitanus* strains encoding the CbbS-mTQ α-carboxysome reporter. In WT, nucleoid staining was homogenous across the cell length and carboxysome foci were confined within the nucleoid signal (PCC = 0.80, n=128 cells) **(Figure 3A)**, consistent with our previous report that the McdAB system uses the nucleoid as a matrix to position carboxysomes **(MacCready et al., 2018)**. In the absence of either Δ*α-mcdA* (PCC = 0.36, n=436 cells) or Δ*α-mcdB* (PCC = 0.33, n=135 cells) α-carboxysome foci were nucleoid excluded at the cell poles **(Figure 3BC)**. Intriguingly, the nucleoids of both the Δ*α-mcdA* and Δ*α-mcdB* strains are shifted away from midcell in the opposing direction of the carboxysome cluster. The data shows that the polar cluster of carboxysomes in *α-mcdAB* mutants are so large and dense that they interfere with gross nucleoid placement in the cell. In some instances, the aggregates grew large enough to locally expand and deform the rod-shaped cell morphology **(Figure 3 – Figure Supplement 1AB)**.

**Figure 3:**
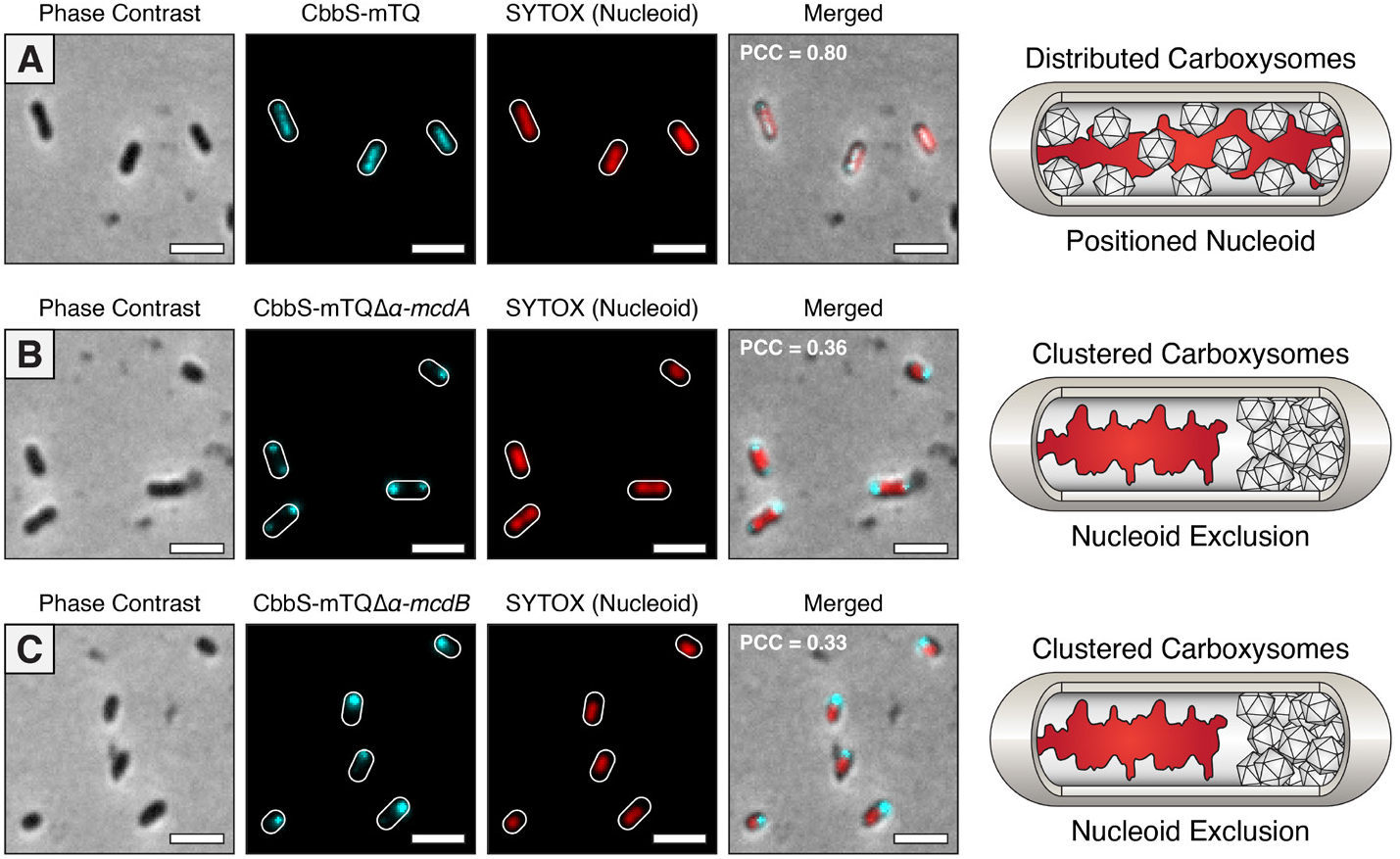
Aggregation of α-carboxysomes leads to nucleoid exclusion. **(A)** CbbS-mTQ signal colocalizes with the nucleoid in WT *S. elongatus*. Aggregated α-carboxysomes are nucleoid excluded in (B) Δα-*mcdA* and (C) Δα-*mcdB* mutants. PCC values calculated from n ≥ 100 cells per cell population. Scale bar = 2 μm.

Upon their discovery, it was frequently noted that the carboxysomes of cyanobacteria and chemoautotrophs are commonly associated with the nucleoid region of the cell **(Jensen and Bowen, 1961; Gantt and Conti, 1969; Shively et al., 1970; Shively et al., 1973; Wolk, 1973).** Our work shows that, for both groups of bacteria, the McdAB system is responsible for this association with the nucleoid. Furthermore, these results suggest that in the absence of the α-McdAB system, nucleoid positioning and cell morphology are altered by the carboxysome cluster. Therefore, in addition to an asymmetric inheritance of carboxysomes in the absence of the McdAB system, the presence of massive carboxysome aggregates has drastic physiological consequences that likely have important implications in the fundamental processes of chromosome segregation, cell division, and bacterial aging.

### α-McdB is targeted to carboxysomes and interacts with α-McdA

We recently showed that β-McdB acts as an adaptor between β-McdA on the nucleoid and β-carboxysomes in the cyanobacterium *S. elongatus*. Bacterial Two Hybrid (B2H) analysis showed that β-McdB associates with individual β-carboxysomes through multiple shell protein interactions **(MacCready et al., 2018)**. To determine whether *H. neapolitanus* α-McdB is similarly targeted to α-carboxysomes, we expressed a second copy of α-McdB as a fluorescent fusion to the fluorescent protein mNeonGreen (mNG) **(Shaner et al., 2013)**, producing mNG-α-McdB, in our native CbbS-mTQ strain. Consistent with mNG-β-McdB loading onto β-carboxysomes in *S. elongatus* **(MacCready et al., 2018)**, we found that mNG-α-McdB colocalized with the CbbS-mTQ signal of *H. neapolitanus* α-carboxysomes (PCC = 0.81; n=173 cells) **(Figure 4A and Video 2)**. Due to the diffuse nature of the α-carboxysome fluorescence signal, as described above, we took advantage of our finding that α-carboxysomes cluster to form bright and clearly resolved puncta at the cell pole without α-McdA. In our *α-mcdA* deletion strain, mNG-α-McdB clearly colocalized with α-carboxysome aggregates (PCC = 0.90; n=355 cells) **(Figure 4B)**. The data not only shows that α-McdB associates with α-carboxysomes, it shows that α-McdA is not required for this association.

**Figure 4:**
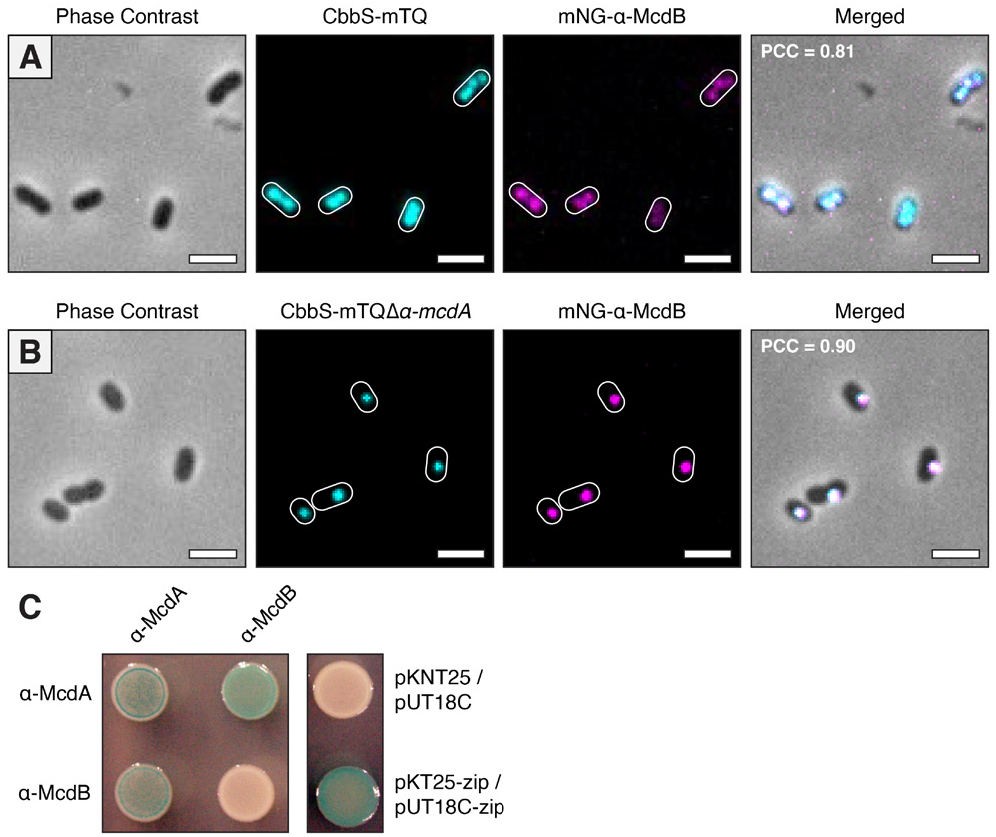
α-McdB loads onto α-carboxysomes and interacts with α-McdA. **(A)** mNG-α-McdB and the carboxysome reporter - CbbS-mTQ – colocalize in WT *S. elongatus*. (B) In the absence of α-McdA, α-McdB colocalizes with carboxysome aggregates. PCC values calculated from n ≥ 100 cells per cell population. Scale bar = 2 μm. (C) Bacterial-2-Hybrid (B2H) of α-McdA and α-McdB. Image representative of 3 independent trials.

Finally, as we did for the β-McdAB system in *S. elongatus*, we sought to determine if α-McdA and α-McdB of *H. neapolitanus* self-associate and directly interact with each other by performing a B2H assay in *E. coli*. Consistent with ParA-family members forming ATP-bound sandwich dimers **(Schumacher, 2007)**, as recently shown for a β-McdA homolog **(Schumacher et al, 2019)**, α-McdA was positive for self-association **(Figure 4C)**. Also, like the β-McdAB system in *S. elongatus*, α-McdA directly interacts with α-McdB. Surprisingly however, we found that *H. neapolitanus* α-McdB did not self-associate in our B2H assay, where we previously found that *S. elongatus* β-McdB strongly self-associates **(MacCready et al., 2018)**. This difference in α- and β-McdB self-association has implications for understanding how McdB proteins are recruited to structurally distinct α- and β-carboxysomes and interact with their cognate McdA ATPase on the nucleoid.

### Defining the conserved features of α-McdAB proteins for bioinformatic searches

We recently found that the□β-McdAB system is not restricted to *S. elongatus*, but rather widespread across β-cyanobacteria, which possess β-carboxysomes (**MacCready et al., 2020**). A surprising finding in this search was that McdAB systems were absent in α-cyanobacteria, which possess α-carboxysomes. Thus, our finding here of an α-McdAB system that positions α-carboxysomes in *H. neapolitanus* was unexpected. We next sought to determine how widespread this system was among α-carboxysome-containing proteobacteria. Given the large range of diversity we previously found among cyanobacterial β-McdA, and especially β-McdB, we approached searching for α-McdA/B outside of *H. neapolitanus* by first performing neighborhood analyses within and around proteobacterial α-carboxysome operons. Using BlastP, we identified α-carboxysome containing proteobacteria within NCBI and JGI IMG databases using α-carboxysome components CsoS2, CsoS4A, or CsoS4B as queries. Next, we searched around each α-carboxysome operon in these organisms to identify *α-mcdAB-like* genes. We classified positive hits for *α-mcdA* as proteins containing the deviant-Walker-A box, which defines the ParA family of ATPases **(Koonin, 2003)**, as well as Walker A’, and Walker B regions. We classified positive hits for *α-mcdB* as a small coding sequence immediately following the *α-mcdA* gene **(MacCready et al., 2020)**.

Alignment of the amino acid sequences of all α-McdA proteins identified through our analyses revealed a high percentage of similarity (~53% pairwise identity), but also revealed novel regions of conservation not present among classical ParA-type proteins **(Figure 5A and Figure 5 - Figure Supplement 1)**. As we found previously in our search for β-McdAB systems, α-McdA proteins were largely conserved and only differed in certain regions, while α-McdB proteins on the other hand displayed extreme diversity. All α-McdB proteins identified shared many of the features present in cyanobacterial β-McdB proteins, including: (i) a well-conserved N-terminally charged region, (ii) an invariant C-terminal tryptophan residue (all ended with the amino acid sequence WPD), (iii) largely polar, (iv) biased amino acid composition, and (v) intrinsic disorder **(Figure 5B and Figure 5 - Figure Supplement 2)**.

**Figure 5:**
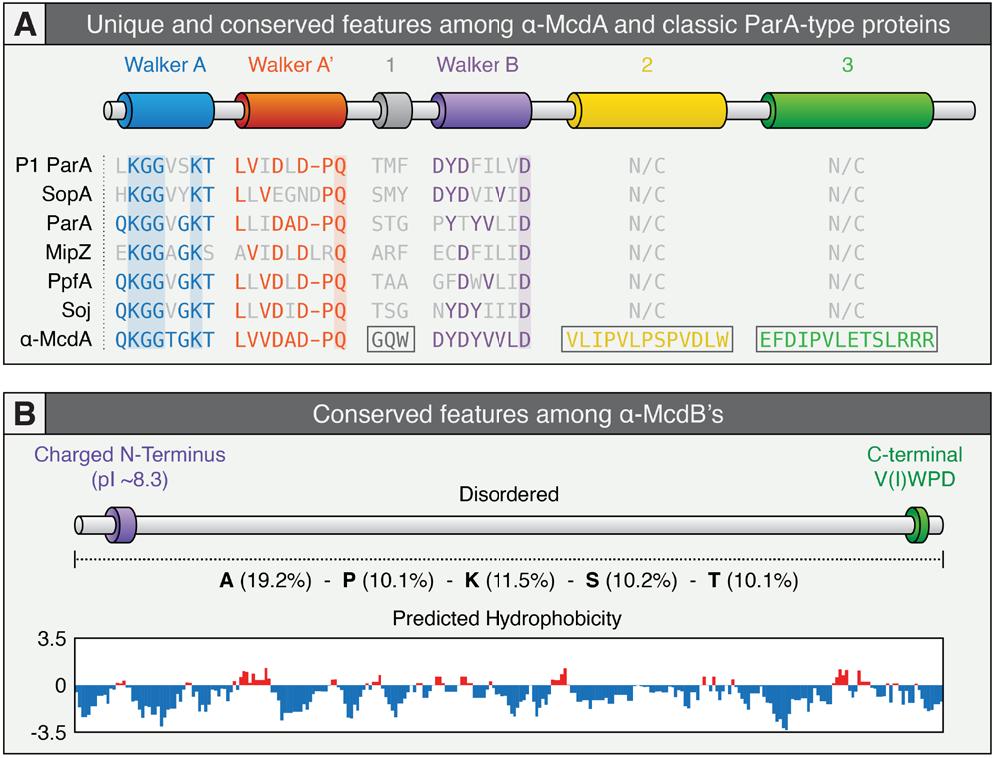
Conserved features among α-McdAB proteins within *cso* operons. **(A)** Features conserved or distinct among identified α-McdA proteins found within the *cso* operon of α-carboxysome containing proteobacteria compared to classical ParA-type proteins involved in the positioning of diverse cargoes. The deviant-Walker A (blue), A’ (red), and B (purple) boxes, are conserved among all ParA family proteins. ParA-type proteins shown: *Escherichia coli*phage P1 ParA (plasmid partitioning—YP_006528), *Escherichia coli* (strain K12) F plasmid SopA (plasmid partitioning — NP_061425), *Caulobacter crescentus* ParA (chromosome segregation—AAB51267), *Caulobacter crescentus* MipZ (cell-division positioning — NP_420968), *Rhodobacter sphaeroides* PpfA (chemotaxis distribution—EGJ21499), and *Bacillus subtilis* Soj (chromosome segregation—NP_391977). (B) General features of α-McdB proteins found within the *cso* operon of α-carboxysome containing proteobacteria. Percent composition of the amino acids alanine, proline, lysine, serine, and threonine present to illustrate strong bias for these amino acids (center). All McdB proteins are highly hydrophilic across the entire primary sequence (bottom).

### α-McdAB systems are widespread among α-carboxysome containing proteobacteria

We previously found that many β-cyanobacteria had *mcdA* and *mcdB* genes at distant loci from the operons encoding carboxysome components. Our identification of highly conserved regions that are specific to α-McdA proteins, along with the conserved features among α-McdB proteins, allowed us to re-search proteobacterial genomes to identify more α-McdA and α-McdB genes that did not fall within or neighbor the α-carboxysome operon. Across 250 α-carboxysome containing proteobacterial genomes, we identified 228 α-McdA sequences (~91% of genomes analyzed) and 278 α-McdB sequences (100% of genomes analyzed) **(Figure 6A)**. These results present a similar trend to what we previously found in several β-cyanobacteria, that β-McdB can exist without the presence of β-McdA and that β-McdB proteins greatly varied in length (51 – 169 amino acids) **(Figure 6A)**. Interestingly, we found that many genomes contained a second *α-mcdB* gene. To our knowledge, this is the first example of two cognate ParA partner protein paralogs potentially involved in the same process. In total, we found three possible genomic arrangements of α-*mcdA* and *α-mcdB* genes: (i) *α-mcdA* and *α-mcdB* were both within the α-carboxysome operon **(Figure 6B)**, (ii) only *α-mcdB* was found within the operon **(Figure 6C)**, or (iii) one *α-*mcdB gene was found within the operon, and a second *α-mcdB* gene was present at a distant locus but next to *α-mcdA* **(Figure 6D)**. Consistent with our identification of orphan McdB proteins, we have recently found that McdB in *S. elongatus* plays an important, but currently unclear, role in carboxysome function outside of its role in positioning carboxysomes with McdA **(Rillema et al. 2020)**. Indeed, ParA-partner proteins involved in the positioning of other protein-based cargos, such as chemotaxis clusters **(Roberts et al., 2012; Ringgaard et al., 2013)**, are also key functional components of the cargo itself. However, we note that 76% of genomes without *α-mcdA* were still in contigs or scaffolds, therefore it is still possible that an *α-mcdA* gene is present and next to the additional *α-mcdB* gene in these genomes.

**Figure 6:**
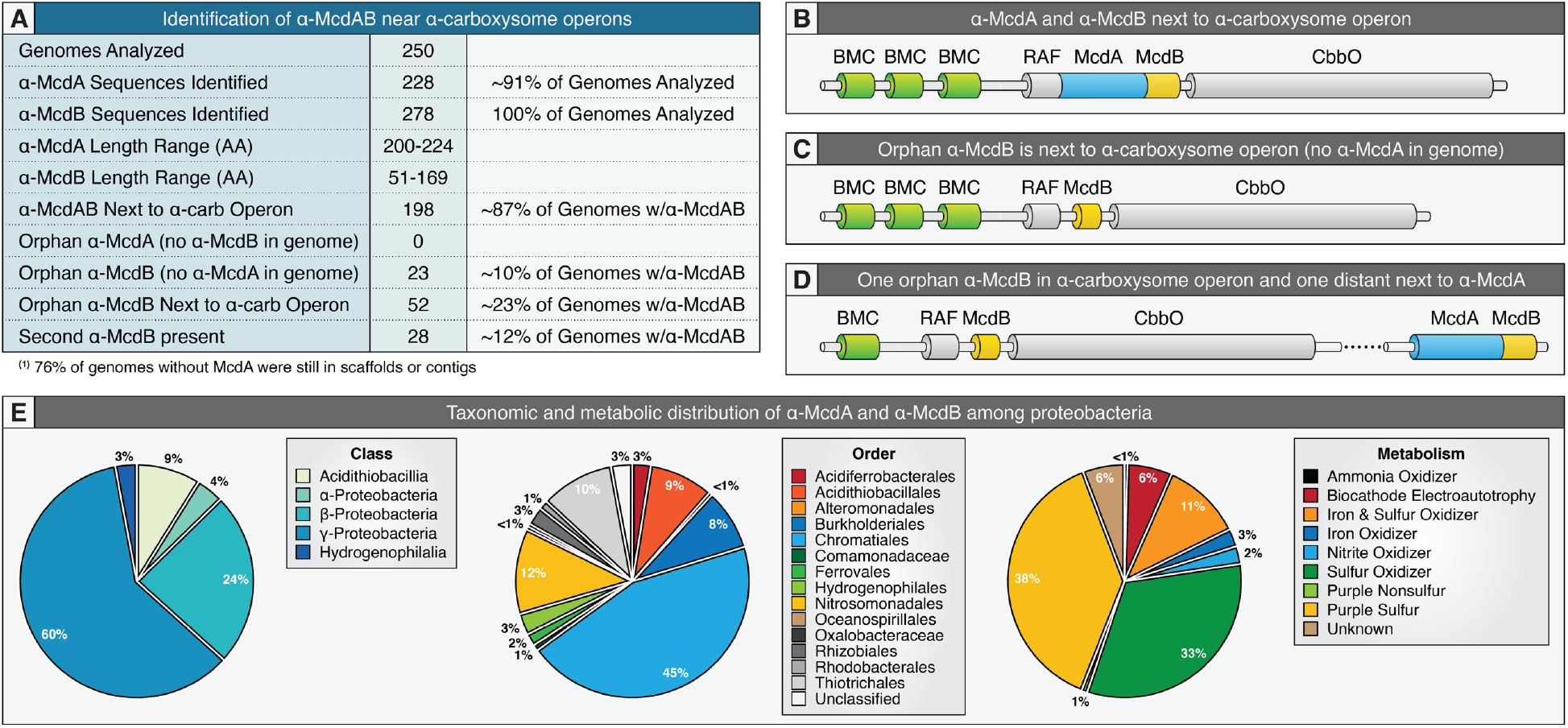
α-McdAB are widespread among α-carboxysome containing proteobacteria. **(A)** Table highlighting the prevalence and genomic context of all identified α-McdAB sequences within α-carboxysome containing proteobacteria. **(B)** Genomic arrangement when α-McdAB are within the *cso* operon. **(C)** Genomic arrangement when only α-McdB is within the *cso* operon. **(D)** Genomic arrangement when one copy of α-McdB is found within the *cso* operon and a second copy of α-McdB is found next to α-McdA at a distant locus. **(E)** α-McdAB are widely distributed among proteobacterial taxonomic classes (left), orders (center) and metabolisms (right).

We also found that α-McdAB systems were present across most major taxonomic classes and orders of proteobacteria; largely present in the class γ-proteobacteria (60% of genomes) and order Chromatiales (~45% of genomes) **(Figure 6E)**. Unlike cyanobacteria, which perform oxygenic photosynthesis, the metabolisms of α-carboxysome containing proteobacteria can greatly vary **(see Figure 1C)**. Despite this, we found that the α-McdAB system was present in nitrite, ammonia, iron, and biocathode utilizers, but largely present in purple sulfur bacteria (38% of genomes), which perform anoxygenic photosynthesis, and sulfur oxidizing chemoautotrophs (33% of genomes) **(Figure 6E)**. Interestingly, the size and quantity of β-carboxysomes in cyanobacteria are directly linked to not only CO_2_ availability but also to light intensity and quality **(Sun et al., 2016; Rohnke et al., 2018; Rohnke et al., 2020)**. Given the diverse metabolic substrates utilized among these α-carboxysome containing proteobacteria, it is possible that α-carboxysome size and quantity are also regulated by nutrient availability in these non-photosynthetic bacteria.

### Cyanobacterial α-carboxysomes likely originated from a proteobacterium lacking α-*mcdA* within their *cso* operon

It is largely believed that α-carboxysomes emerged in proteobacteria and were subsequently horizontally transferred into cyanobacteria, thus creating the distinct phylogenetic clade of α-cyanobacteria **(Badger et al., 2002; Badger and Price, 2003; Marin et al., 2007; Badger and Bek, 2008; Rae et al., 2013)**. However, following our recent study **(MacCready et al., 2020)**, why α-cyanobacteria lack the McdAB system remained an outstanding question. In an attempt to better understand α-carboxysome evolution among α-cyanobacteria and proteobacteria, we constructed a Maximum Likelihood phylogenetic tree inferred using a concatenation of the major α-carboxysome components CbbL, CbbS, CsoS3, CsoS4A, and CsoS4B. We note that the major carboxysome component CsoS2 was omitted from this analysis due to its intrinsically disordered nature **(Oltrogge et al., 2020)**, which has relaxed selection due to the lack of structural constraint.

Recall that we found α-carboxysome containing proteobacteria that encoded two α-McdB genes, where one *α-mcdB* gene is encoded within the *cso* operon without an adjacent α-*mcdA* gene, and the other is encoded next to α-*mcdA* at a distant locus away from the *cso* operon **(see Figure 6D)**. Intriguingly, we found that these proteobacteria all share a common ancestor and possessed α-carboxysomes that were more closely related to α-cyanobacterial α-carboxysomes than to proteobacterial α-carboxysomes that had both α-*mcdA* and *α-mcdB* encoded within the *cso* operon **(Figure 7A)**. Indeed, while *H. neapolitanus* possess both α-*mcdA* and *α-mcdB* within their *cso* operon **(Figure 7B)**, many other proteobacterial species, such as *Thiohalospira halophila* DSM 15071, only possess *α-mcdB* within their *cso* operon; α-*mcdA* and the second *α-mcdB* gene are encoded at a distant locus **(Figure 7C)**. Thus, our phylogenetic tree suggests that α-cyanobacteria, such as *Cyanobium gracile* PCC 6307, likely horizontally inherited a *cso* operon from a proteobacterium that encoded α-*mcdA* and a second *α-mcdB* gene away from the *cso* operon **(Figure 7D)**. While it is possible that α-cyanobacteria inherited the orphan *α-mcdB* gene present within the proteobacterial *cso* operon, it is likely that this protein no longer provided a fitness advantage in the absence of α-McdA, or the second α-McdB protein, and was therefore lost.

**Figure 7:**
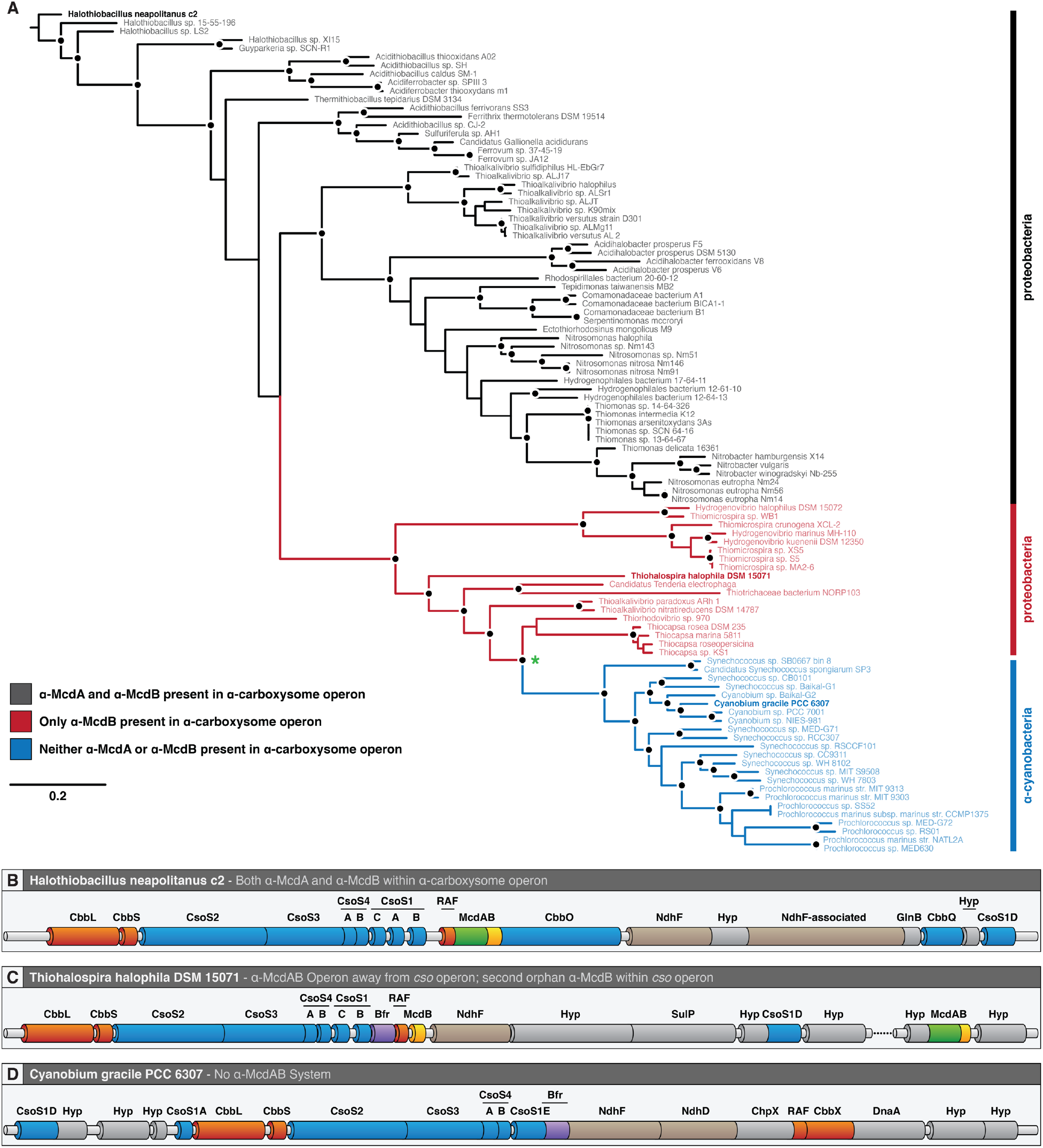
α-carboxysome evolution among proteobacteria and cyanobacteria. **(A)** Inferred phylogeny of α-carboxysome containing proteobacteria and cyanobacteria. Line colors: α-McdAB found within *cso* operon (black), only α-McdB found within *cso* operon (red), and neither α-McdAB found within *cso* operon. Black dot represents >70% bootstrap support (500 replicates). Green asterisk represents a shared common ancestor among α-cyanobacteria and proteobacteria that lack α-McdA in their *cso* operon. (B) Genomic arrangement of the *H. neapolitanus cso* operon. (C) Genomic arrangement of the *T. halophila cso* operon. (D) Genomic arrangement of the *C. gracile cso* operon.

### α-McdAB are distinct from cyanobacterial Type 1 and Type 2 β-McdAB systems

Given the large quantity of α- and β-McdAB systems that we have identified across cyanobacteria and proteobacteria, we next wanted to identify similarities and differences among these subtypes. We previously identified two distinct β-McdAB systems exclusive to β-cyanobacteria, with Type 2 systems being more ancestral and more abundant than Type 1 **(MacCready et al, 2020)**. Here, we found that α-McdA proteins share more features with β-McdA Type 2 proteins, including: (i) the presence of the signature lysine residue in the deviant-Walker A box, which defines the ParA family but is absent in β-McdA Type I, (ii) a conserved central tryptophan residue in region 1, (iii) a highly similar Walker B box, and (iv) greater conservation in region 4 **(Figure 8A)**. This further demonstrates the divergence of Type 1 β-McdA proteins (present in *S. elongatus*), which lack the signature lysine residue in the deviant-Walker A box and possesses a large mid-protein extension **(MacCready et al, 2020)**.

**Figure 8:**
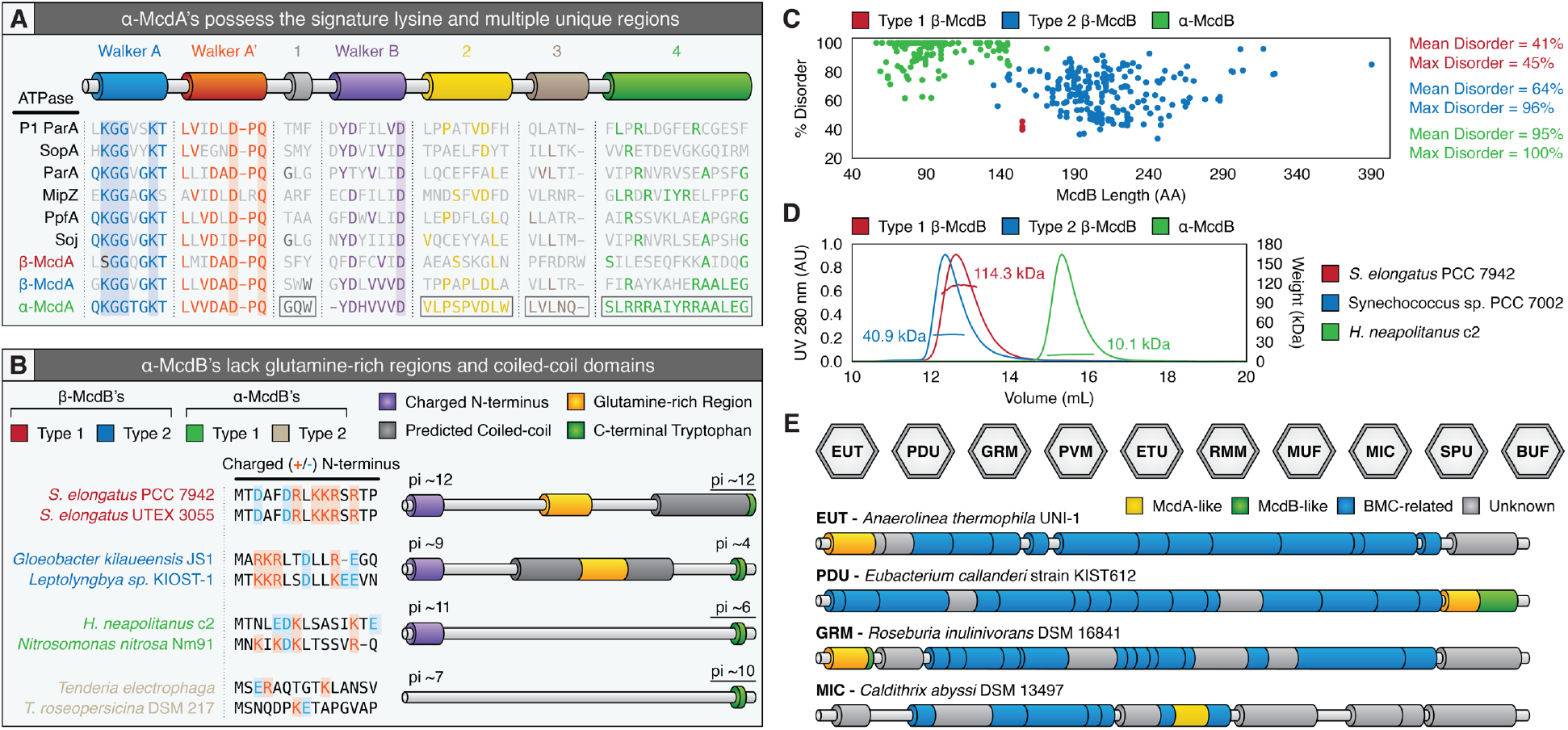
Similarities and differences among all known McdB proteins. **(A)** Features conserved or distinct among α-McdA, β-McdB Type 1, β-Type 2, and diverse classical ParA-type proteins. Known conserved ParA regions, deviant-Walker A (blue), A’ (red), and B (purple) boxes, are conserved among all classic ParA proteins. ParA-type ATPase proteins shown: *Escherichia coli* phage P1 ParA (plasmid partitioning—YP_006528), *Escherichia coli* (strain K12) F plasmid SopA (plasmid partitioning—NP_061425), *Caulobacter crescentus* ParA (chromosome segregation—AAB51267), *Caulobacter crescentus* MipZ (chromosome segregation — NP_420968), *Rhodobacter sphaeroides* PpfA (chemotaxis distribution— EGJ21499), and *Bacillus subtilis* Soj (chromosome segregation—NP_391977). **(B)** α-McdB Type 1 and Type 2 proteins lack predicted central coiled-coil and glutamine-rich regions found in β-McdB Type 1 and Type 2 proteins. α-McdB Type 2 proteins lack a charged N-terminal region. **(C)** PONDR disorder scatter plot for all β-McdB Type 1 (red), β-McdB Type 2 (blue), and α-McdB (green) proteins. **(D)** SEC-MALS plot for a representative β-McdB Type 1 (*S. elongatus* McdB; monomer MW = 17 kDa), β-McdB Type 2 (*Synechococcus sp*. PCC 7002 McdB; monomer MW = 21 kDa), and α-McdB (*H. neapolitanus* McdB; monomer MW = 10 kDa). **(E)** McdA/B-like sequences genomically neighbor BMC components across diverse microbes.

ParB proteins that connect plasmid or chromosomal cargo to their cognate ParA ATPase use a charged N-terminal region to interact with ParA and stimulate its ATPase activity **(Baxter and Funnell, 2014; Badrinarayanan et al., 2015)**. We found that all McdB types encoded near their cognate McdA also possess conserved highly charged N-termini **(Figure 8B)**. But in proteobacteria where the second *α-mcdB* gene is orphaned in the *cso* operon, this highly charged N-terminus is absent **(Figure 8B)**. Given this observation, we designate α-McdB proteins with the charged N-terminal region next to α-McdA proteins as Type 1, and α-McdB proteins lacking the charged N-terminus and orphaned in the *cso* operon as Type 2. Further dividing α- and β-McdB proteins, we find that α-McdB proteins lack the predicted coiled-coil region and mid-protein glutamine-rich stretches conserved among β-McdB proteins **(Figure 8B)**. Despite these differences among subtypes, all McdB proteins possess the invariant C-terminal tryptophan residue.

Our finding that many proteobacteria possessed an additional distinct copy of α-McdB was intriguing; we have yet to identify a cyanobacterial species that possesses two *mcdB* genes **(MacCready et al., 2020)**. The need for two α-McdB proteins is not clear. However, given the lack of a charged N-terminus, which has been shown in several ParA-based positioning systems to be responsible for interacting with their cognate ParA, these paralogs likely play a role in modulating interactions with α-McdA and possibly tuning the stimulation of McdA ATPase activity. For instance, one α-McdB protein could interact with α-McdA and the other paralog may buffer this association. The interaction between two paralogs to regulate function is not uncommon. One such example is the activity of FtsZ paralogs (FtsZ1 and FtsZ2) involved in archaeal cell division **(Liao et al., 2020)** and chloroplast division **(Reviewed in: Chen et al., 2018)**. In the case of chloroplasts, these paralogs self-assemble into heteropolymers that further assemble into a ring-like structure at mid-chloroplast to define the site of binary fission. FtsZ1 increases FtsZ2 turnover, which is thought to facilitate FtsZ2 remodeling during chloroplast constriction **(Terbush and Osteryoung, 2012)**. In this way, FtsZ1 functions as a regulator of FtsZ2 activity and is a highly conserved feature throughout photosynthetic lineages **(Terbush et al., 2018)**. Elucidating how α-McdB paralogs influence carboxysome function and organization with α-McdA is an exciting avenue of future research.

A conserved feature across all McdB types we identified is intrinsic disorder, but to varying degrees **(Figure 8C)**. While β-McdB Type 1 proteins are on average 41% disordered and β-McdB Type 2 proteins are on average 64% disordered, α-McdB proteins are significantly more disordered (~95% disordered) **(Figure 8C)**. This dramatic difference in predicted disorder for α-McdB proteins is in part likely due to the lack of the predicted central coiled-coil region present in both β-McdB types **(see Figure 8B)**. Given the large differences in predicted disorder and predicted coiled-coil regions among α- and β-McdB proteins, we performed Size Exclusion Chromatography - Multiple Angle Laser Light Scattering (SEC-MALS) to determine the oligomeric state of purified α- and β-McdB proteins. We found that the Type 1 β-McdB of *S. elongatus* formed a hexamer, *Synechococcus sp*. PCC 7002 Type 2 β-McdB formed a dimer, and consistent with lacking a predicted coiled-coil, *H. neapolitanus* α-McdB remained monomeric **(Figure 8D)**. This finding explains our B2H data, where β-McdB of *S. elongatus* was found to self-associate strongly, whereas *H. neapolitanus* α-McdB showed no self-association at all **(Figure 4C)**. These results suggest the predicted coiled-coil domains exclusive to β-McdB proteins are required for oligomerization and are possibly involved in the function of McdB proteins in their respective carbon-fixing bacterium.

Lastly, an outstanding question is whether the McdAB system is restricted to carboxysome BMCs. While carboxysomes are the only known anabolic BMC, several other catabolic BMCs exist, including: (I) EUT (<u>E</u>thanolamine <u>UT</u>ilization microcompartment), (II) PDU (1,2-<u>P</u>ropanediol <u>U</u>tilization microcompartment), (III) GRM (<u>G</u>lycyl <u>R</u>adical enzyme-containing <u>M</u>icrocompartment), (IV) PVM (<u>P</u>lanctomycetes and <u>V</u>errucomicrobia <u>M</u>icrocompartment), (V) ETU (<u>E</u>Thanol-<u>U</u>tilizing microcompartment), (VI) RMM (<u>R</u>hodococcus and <u>M</u>ycobacterium <u>M</u>icrocompartment), (VII) MUF (<u>M</u>etabolosome of <u>U</u>nknown <u>F</u>unction microcompartment), (VIII) MIC (<u>M</u>etabolosome with <u>I</u>ncomplete <u>C</u>ore microcompartment), (VIIII) SPU (<u>S</u>ugar <u>P</u>hosphate <u>U</u>tilizing microcompartment), and (X) BUF (<u>B</u>MC of <u>U</u>nknown <u>F</u>unction) **(Figure 8E)**. In four such examples, we find McdB- and/or McdA-like sequences within or neighboring these BMC operons **(Figure 8E)**. In cases where McdB-like sequences are observed, they all possess a C-terminal aromatic residue (tyrosine instead of tryptophan) within the last 4 amino acids, a feature that is invariant across all carboxysome-associated McdB proteins we have identified to date. Moreover, these putative McdB proteins end with the residues LL; ~8% of β-McdB proteins end with LL and ~28% end with LLD. It is intriguing that many proteins involved in the assembly of viral- or phage-capsids also encode for an aromatic residue (tryptophan) at their C-terminus **(Deeb, 1973; Skoging and Liljeström, 1998; Tsuboi et al. 2003; Komla-Soukha and Sureau 2006; Marintcheva et al. 2006; Johnson et al., 2020)**. Given the capsid-like icosahedral structure of BMCs, it is attractive to speculate that this C-terminal aromatic residue may play a role in McdB association with their cognate BMC.

Collectively, our results show that α-McdAB systems are widespread among α-carboxysome containing proteobacteria and function as an anti-aggregation mechanism to ensure proper distribution and inheritance of α-carboxysomes following cell division. Our results have important implications for understanding the evolution and function of diverse McdA and McdB proteins as it relates to structurally distinct α- and β-carboxysomes, and also have much broader implications for understanding the equidistant positioning of diverse catabolic BMCs across the domain of bacteria.

## Acknowledgements

We would like to thank Dr. JK Nandakumar for assistance in the SEC-MALS experiments. This work was supported by the National Science Foundation to A.G.V. (Award No. 1817478 and CAREER Award No. 1941966), NSF GRP Award to L.T. (DGE 1841052), Rackham Graduate Student Research Grant to L.T., and from research initiation funds provided by the MCDB Department to A.G.V., University of Michigan and the Michigan Life Sciences Fellows Program to J.S.M. We’d also like to thank Jeffrey Harrison from the UM Microscopy Core for assistance with TEM preparation and imaging.

## Supplemental Information

**Supplemental Figure 1:**
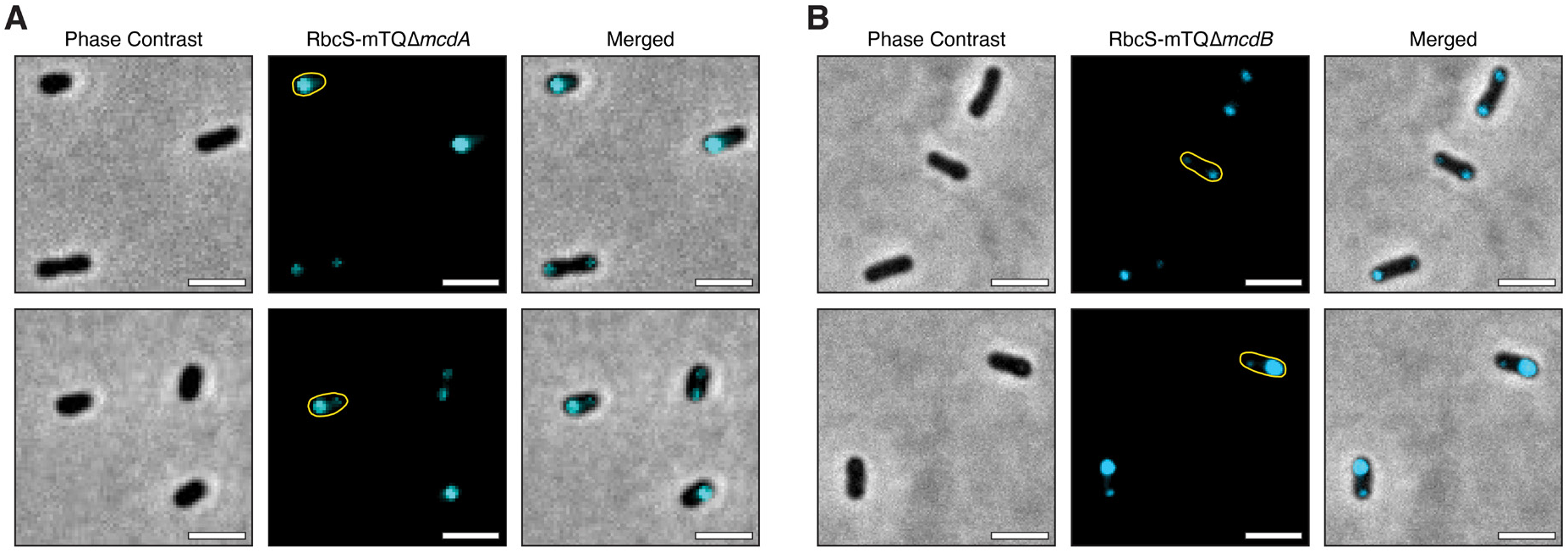
Aggregation of α-carboxysomes caused local bulging in cellular rod-shape morphology in the absence of **(A)** α-McdA, or McdB. Scale bar = 2 μm.

**Supplemental Figure 2:**
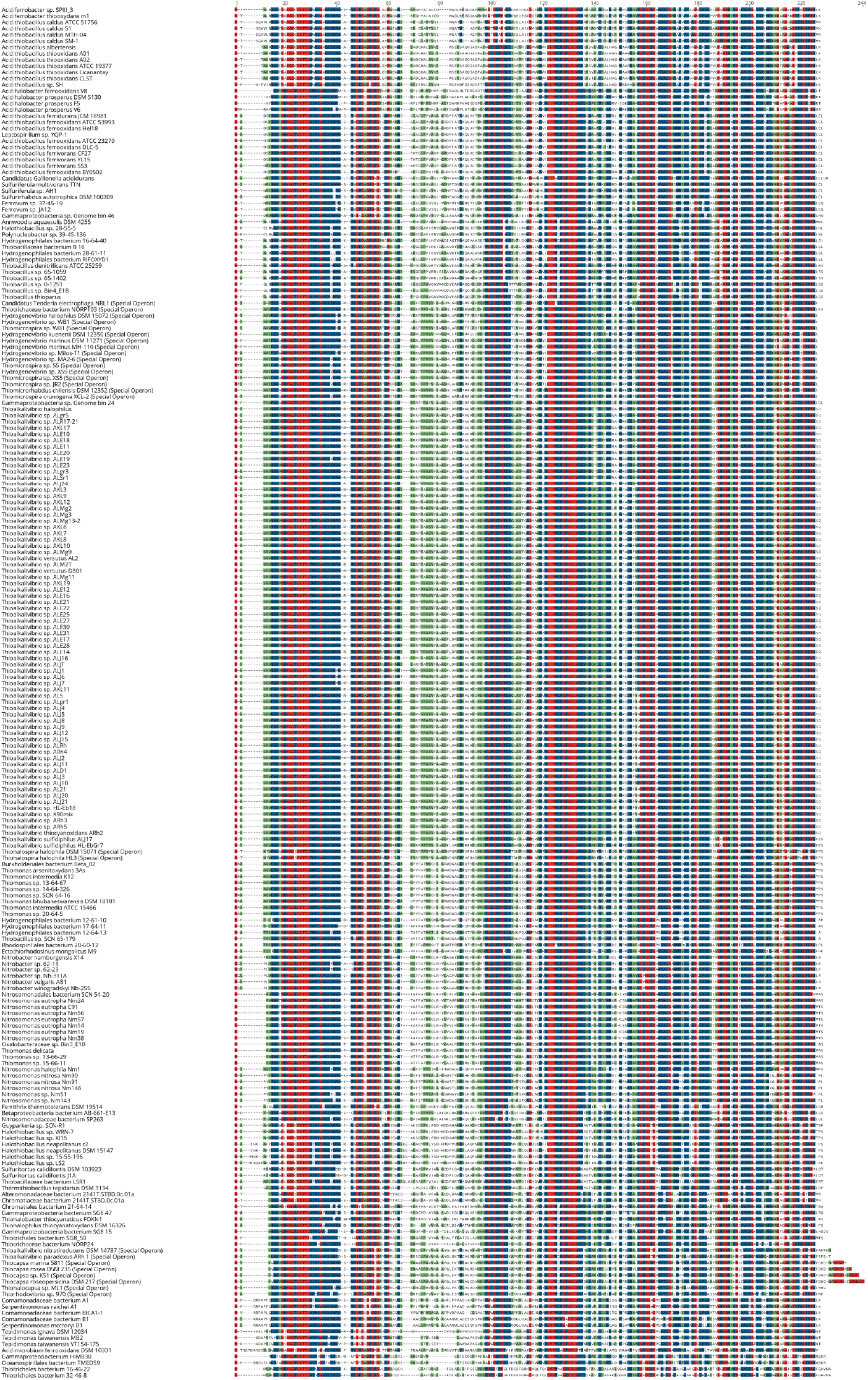
Multiple sequence alignment of identified α-McdA sequences from proteobacteria

**Supplemental Figure 3:**
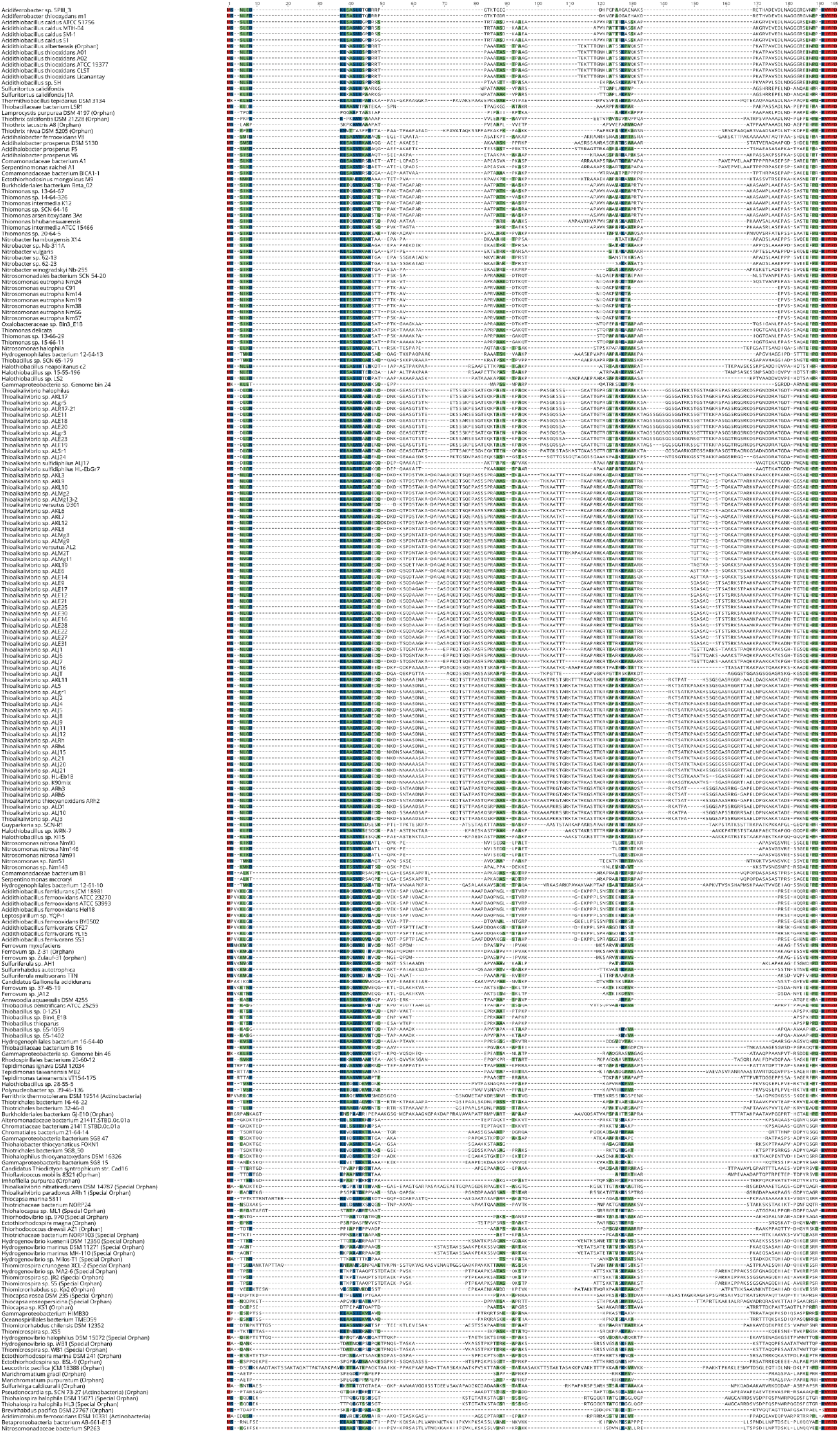
Multiple sequence alignment of identified α-McdB sequences from proteobacteria

## Methods

### Strains and growth conditions

All mutants described in this study were constructed with wild-type *H. neapolitanus* c2. Cultures were grown in S6 medium and incubated shaken at 130 RPM, at 30°C, in air supplemented with 5% CO_2_. Strains are preserved frozen at −80°C in 10% DMSO.

### Construct designs and cloning

All constructs were generated using Gibson Assembly and verified by sequencing. Fragments for assembly were synthesized by PCR or ordered as a gBlock (IDT). Constructs contained flanking DNA that ranged from 750 to 1100 bp in length upstream and downstream of the targeted insertion site to promote homologous recombination into target genomic loci. Cloning of plasmids was performed in chemically competent *E. coli* Top10 or Stellar cells (Takara Bio).

### Transformation in *H. neapolitanus* c2

Competent cells of *H. neapolitanus* were generated by growing 1 L of log culture in 2.8 L baffled flasks. Cultures were harvested by centrifugation at 3,000xg for 45 minutes. Pellets were resuspended and washed twice with 0.5 volumes of ice-cold millipore water. The resulting pellet was resuspended in 1×10-^3^ volumes of ice-cold millipore water. These competent cells were used immediately or frozen at −80°C for future use. Frozen competent cells were thawed for 4 hr or overnight at 4°C. Competent cells were mixed with 5 μL plasmid DNA (1.5-5 μg) and incubated on ice for 5 minutes. This mixture is then transferred to a tube containing 5 mL ice-cold S6 medium without antibiotics and incubated on ice for 5 minutes. Transformations were recovered for 16-36 hours, shaken at 130 RPM, at 30°C, in air supplemented with 5% CO_2_. Clones were selected by plating on selective medium with antibiotics. Colonies were restreaked. Restreaked colonies were verified for mutation by PCR.

### Native fluorescent fusions

For native CbbS fluorescent fusion, the fluorescent protein mTurquoise (mTq) was attached to the 3’ region of the native *cbbS* coding sequence, separated by a GSGSGS linker. A kanamycin resistance cassette was inserted immediately downstream of the *cbbS* coding sequence. The mutant was selected by plating on S6 agar plates supplemented with 50 μg/mL kanamycin. The mutation was verified by PCR.

### Single gene deletions

*mcdA* (Hn0912) and *mcdB* (Hn0911) are the second and third genes in their operon. The deletion construct of mcdA (Hn0912) was created by replacing the mcdA coding sequence with a spectinomycin resistance cassette. The promoter upstream of Hn0913 was duplicated and inserted downstream of the antibiotic cassette to minimize disruption of downstream genes. The deletion construct of *mcdB* was created by removing *mcdB* (Hn0911), along with the 2 upstream genes (Hn0913 and Hn0912). These genes were replaced with a spectinomycin resistance cassette, followed by the promoter upstream of Hn0913 and the codon-optimized genes Hn0913-Hn0912. This allowed for a clean deletion of the *mcdB* coding sequence. Mutants were selected by plating on S6 agar plates supplemented with 50 μg/mL spectinomycin. The mutation was verified by PCR.

### Exogenous expression of mNG-McdB

The fluorescent protein mNeonGreen (mNG) was attached to the 5’ region of the *mcdB* coding sequence, separated by a GSGSGS linker. Gene was codon-optimized, placed under the expression of a P_trc_ promoter, and inserted into a neutral site, located between genes Hn0933 and Hn0934. Mutants were selected by plating on S6 agar plates supplemented with 25 μg/mL chloramphenicol. The mutation was verified by PCR.

### Nucleoid staining

Cells were harvested by centrifugation at 5,000xg for 5 minutes. Following centrifugation, the cells were washed in PBS. The resulting cell pellet was resuspended in 50 μL PBS and stained with SytoxOrange at a final concentration of 25 μM for 5-10 minutes immediately prior to imaging.

### Fluorescence imaging

All live-cell microscopy was performed using exponentially growing cells. 3 μL of cells were dropped onto a square of 2% agarose + S6 pad and imaged on a glass-bottom dish (MatTek Life Sciences). Exogenous expression mutants were not induced. All images obtained resulted from leaky expression of the P_trc_ promoter. Exogenous expression mutants were compared back to an empty vector control. For carboxysome movies, images were taken at an interval of 1 minute for a duration of 15 minutes.

### Transmission Electron Microscopy

Log cultures of *H. neapolitanus* c2 were pelleted and fixed overnight at 4°C in 2.5% glutaraldehyde in 0.1 M Sorensen’s buffer (pH 7.4). After several washes in 0.1M Sorensen’s buffer, cells were post-fixed overnight at 4°C in 1% osmium tetraoxide in 0.1 M Sorensen’s buffer. Cells were washed until the solution became clear. Cells were then immobilized in 1% agarose and sliced thinly into 0.5 mm slices. Slices were dehydrated in an increasing series of ethanol or acetone (30-100%) for 10 minutes each. Slices were infiltrated with Embed812 resin (25% increments for 1-16 hours each at room temperature). A final incubation in full strength resin was performed at room temperature under vacuum. Cells were then embedded in blocks, incubated under vacuum overnight, and transferred to a 60°C oven to polymerize for 1-2 days. Thin sections of approximately 50 nm were obtained by using a Leica UC7 Ultramicrotome. Sections were post-stained with 7% uranyl acetate for 10 minutes, Reynold’s lead citrate for 5 minutes. and visualized on an JEOL 1400-plus transmission electron microscope equipped with a XR401 AMT sCMOS camera.

### Bacterial Two-Hybrid

N-terminal T18 and T25 fusions of McdA and McdB were constructed using plasmid pKT25, pKNT25, pUT18C and pUT18, sequence-verified and co-transformed into E. coli BTH101 (Karimova et al., 1998). Several colonies of T18/T25 cotransformants were isolated and grown in LB medium with 100 μg/ml ampicillin, 50 μg/ml kanamycin and 0.5 mM IPTG overnight at 30°C with 225 rpm shaking. Overnight cultures were spotted on indicator Xgal plates supplemented with 100 μg/ml ampicillin, 50 μg/ml kanamycin and 0.5 mM IPTG. Plates were incubated at 30°C up to 48 hr before imaging.

### α-McdAB Homolog Search and Neighborhood Analysis

Identification of all α-carboxysome containing proteobacteria within the NCBI and JGI databases was performed via BlastP using the α-carboxysome components CsoS2, CsoS4A, or CsoS4B as queries. Neighborhood analyses for McdA- and McdB-like sequences were then carried out within these resulting genomes by manually searching 4000 bp upstream and downstream of the *cso* operon. Multiple sequence alignment for identified putative McdA proteins was performed using MAFFT 1.3.7 under the G-INS-I algorithm, whereas the E-INS-I algorithm was used for McdB proteins due to long gaps caused by intrinsic disorder. Subsequent identification of McdA- and McdB-like sequences away from the *cso* operon was performed via BlastP using highly conserved regions/features identified from the multiple sequence alignments.

### α-McdB Sequence Analysis

Coiled-coil predictions for α-McdB proteins were performed using DeepCoil **(Ludwiczak et al. 2019)**. α-McdB protein disorder predictions were conducted using PONDR with the VL-XT algorithm **(Romero et al. 1997, 2001; Li et al. 1999)**. Analysis of α-McdB hydrophobicity was performed using ProtScale using the Kyte and Doolittle scale **(Kyte and Doolittle 1982; Gasteiger et al. 2005)**.

### Phylogenetic Inference

Ortholog sequences for *H. neapolitanus* CbbL (Hneap_0922), CbbS (Hneap_0921), CsoS3 (Hneap_0919), CsoS4A (Hneap_0918), and CsoS4B (Hneap_0917) were obtained via BLASTp for each proteobacterium and cyanobacterium. Multiple sequence alignments for each protein sequence were performed using MAFFT 1.3.7 under the G-INS-I algorithm and BLOSUM62 scoring matrix. The five resulting alignments were then concatenated into one alignment using Geneious 11.1.5. Regions of low conservation within the resulting alignment were removed using gBlocks 0.91 b **(Castresana 2000; Talavera and Castresana 2007)**. A phylogenetic tree was then estimated with maximum likelihood analyses using RAxML 8.2.11 under the LG+Gamma scoring model of amino acid substitution. Bootstrap values were calculated from 500 replicates.

### Expression and Purification of McdB Homologs

McdBs were expressed with an N-terminal His-SUMO tag off a pET11b plasmid. Vectors were transformed into competent BL21-AI cells. Expression cultures were grown in 2 L of LB + carbenicillin (100 μg/mL) at 37 ^o^C to an OD600 of 0.6. Cultures were then induced with final concentrations of IPTG at 1 mM and L-arabinose at 0.2% (w/v) and allowed to grow for 4 hrs. at 37 ^o^C. Cultures were pelleted and resuspended in 80 mL lysis buffer (50 mM Tris-HCl pH 8.5, 300 mM KCl, 5 mM BME, 0.05 mg/mL lysozyme, 0.05 μL/mL benzonase, protease inhibitor) on ice. Cells were lysed using a tip sonicator at 50% power with 10/20 sec. on/off cycles for 6 mins at 4 ^o^C. Lysates were spun down at 35,000 x g at 4 ^o^C, and supernatants loaded onto 5 mL HP HIS-TRAP columns (GE Healthcare Life Sciences) equilibrated in buffer A (50 mM Tris-HCl pH 8.5, 300 mM KCl, 5 mM BME, 20 mM imidazole). Proteins were eluted using a 5-100% gradient of buffer B (50 mM Tris-HCl pH 8.5, 300 mM KCl, 5 mM BME, 500 mM imidazole) via an AKTA pure system (GE Healthcare Life Sciences). Peak fractions were pooled and diluted in buffer A to a final imidazole concentration < 100 mM. 300 μL of purified His-Ulp1 was added and reactions were incubated at 30 ^o^C to cleave the His-SUMO tag. Reactions were then passed over 5 mL HP HIS-TRAP columns to remove tags and enzymes. Flow-through was concentrated and passed over a SEC column (HiLoad 16/600 Superdex 200 pg; GE Healthcare Life Sciences) equilibrated in buffer C (50 mM Tris-HCl pH 8.5, 150 mM KCl, 5 mM BME, 10% glycerol) using an AKTA pure system. Peak fractions were pooled and stored at −80°C.

### Size-Exclusion Chromatography with Multi-Angle Light Scattering (SEC–MALS)

For each McdB homolog analyzed, 500 μL of sample at 1.5 mg/mL was passed over a SEC column (Superdex 200 Increase 10/300 GL; GE Healthcare Life Sciences) at a flow rate of 0.15 mL/min in buffer (50 mM Tris-HCl pH 8.5, 150 mM KCl, 5 mM BME) at 4 °C. Following SEC, the samples were analyzed using an A280 UV detector (AKTA pure; GE Healthcare Life Sciences), the DAWN HELEOS-II MALS detector (Wyatt Technology) and the OptiLab rEX refractive index detector (Wyatt Technology). The data were analyzed to calculate mass using ASTRA software (Wyatt Technology). Bovine serum albumin was used as the standard for calibration.

